# Evidence of recent increased pathogenicity within the Australian *Ascochyta rabiei* population

**DOI:** 10.1101/2020.06.28.175653

**Authors:** P Sambasivam, Y Mehmood, I Bar, J Davidson, K Hobson, K Moore, R Ford

**Affiliations:** Environmental Futures Research Institute, School of Environment and Science, Griffith University, QLD, 4111 Australia; South Australian Research and Development Institute, Hartley Grove, Urrbrae SA 5064; Department of Primary Industries Tamworth Agricultural Institute, Calala, NSW 2340 Australia

## Abstract

Ascochyta Blight (AB), caused by *Ascochyta rabiei* (syn *Phoma rabiei*), is the major endemic foliar fungal disease affecting the Australian chickpea industry, resulting with potential crop loss and management costs. This study was conducted to better understand the risk posed by the Australian *A. rabiei* population to current resistance sources and to provide informed decision support for chemical control strategies. Recent changes in the pathogenicity of the population were proposed based on disease severity and histopathological observations on a host set. Controlled environment disease screening of 201 isolates on the host set revealed distinct pathogenicity groups, with 41% of all isolates assessed as highly aggressive and a significant increase in the proportion of isolates able to cause severe damage on resistant and moderately resistant cultivars since 2013. In particular, the frequency of highly aggressive isolates on the widely adopted PBA HatTrick cultivar rose from 18% in 2013 to 68% in 2017. In addition, isolates collected since 2016 caused severe disease on Genesis 090, another widely adopted moderately resistant cultivar and on ICC3996, a commonly used resistance source. Of immediate concern was the 10% of highly aggressive isolates able to severely damage the recently released resistant cultivar PBA Seamer (2016). Histopathology studies revealed that the most aggressive isolates were able to germinate, develop appressoria and invade directly through the epidermis faster than lower aggressive isolates on all hosts assessed, including ICC3996. The fungal invasion triggered a common reactive oxygen species (ROS) and hypersensitive response (HR) on all assessed resistant genotypes with initial biochemical and subsequent structural defence responses initiated within 24 hours of inoculation by the most highly aggressive isolates. These responses were much faster on the less resistant and fastest on the susceptible check host, indicating that speed of recognition was correlated with resistance rating. This will inform fungicide application timing so that infected crops are sprayed with prophylactic chemistries prior to invasion and with systemic chemistries after the pathogen has invaded.

## Introduction

Chickpea (*Cicer arietinum* L.) is an important staple pulse crop grown and consumed globally as a source of dietary protein, particularly in central Asia and Africa (Gan et al., 2006, Harveson et al., 2011, Kanouni et al., 2011). Australia is the second largest producer of chickpea (FAO 2018), however, yield and quality is significantly affected by Ascochyta Blight (AB) caused by the necrotrophic fugal pathogen, *Ascochyta rabiei* (Pass.) Labr. (syn. *Phoma rabiei*) (Taylor and Ford, 2007; Atik et al., 2011; Bayaa and Chen, 2011; Moore et al., 2015). The disease occurs in major chickpea growing regions and reduces yield, seed quality and limits production worldwide (Nene, 1982; Kaiser et al., 2000; Siddique et al., 2000; Chongo et al., 2003; Millan et al., 2003; Sarwar et al., 2012; Sharma and Ghosh, 2016).

The disease may affect all above ground plant tissues and occur at any growth stage depending on favourable environmental conditions (Armstrong-Cho et al., 2001). Damage from lesions on leaves, stems and pods reduces photosynthesis with subsequent stem breakage, pod abortion, seed staining and even plant death (Pande et al., 2005). Management is based on growing chickpea varieties with improved resistance together with a fungicide regime. Fungicide is applied usually under a multiple spray regime and prior to predicted rainfall events to prevent or attempt to manage epidemics (Moore et al., 2016; Pulse Australia, 2018).

Genetic variation and concomitant adaptation within *A. rabiei* populations has led to the erosion of resistance (Peever et al., 2012; Mahiout et al., 2015; Vafaei et al., 2015; Mehmood et al., 2017; Tekin et al., 2017). This has created highly aggressive isolates that are able to cause significant disease on commonly deployed cultivars (Varshney et al., 2007; Mehmood et al., 2017). Selection for stable and durable resistance to prevailing isolates within a growing region, requires an understanding of the pathogenic structure in the population and identification of isolates for deployment in future resistance breeding strategies. Also, knowledge of the interaction between highly aggressive isolates and the commonly grown cultivars should contribute to improved choice and timing of fungicide application.

Whilst breeders have moved away from single source resistance to employing pyramided polygenic resistance (Thabuis et al., 2004; Reeves et al., 2013), longevity of newly deployed “ resistant” cultivars remains dependent on the speed of isolate evolution and population adaptation (Moore et al., 2015; Mehmood et al., 2017). Accordingly, to assess this risk, variation in isolate aggressiveness has been assessed in many global regions including India (Vir and Grewal, 1974), the Palouse region of USA (Chen et al., 2004) Spain (Navas-Cortes et al., 1998), Pakistan (Jamil et al., 2000; Iqbal et al., 2004; Ali et al., 2012), Canada (Chongo et al., 2004), Syria (Atik et al., 2013; Imtiaz et al., 2015) and Australia (Mehmood et al., 2017). Within these studies, groups of isolates that interact in a differential manner with a host genotype set have been identified possessing a range of aggressiveness within the populations. Where specific discreet factors related to host interaction were noted within a range of isolate pathogenicity, pathotypes were proposed (Vir and Grewal, 1974; Udupa et al., 1998; Atik et al., 2013; Imtiaz et al., 2015; Baite and Dubey, 2018). In Syria, Udupa et al (1998) originally concluded the existence of three pathotypes: pathotype 1 (low aggressive), pathotype II (aggressive) and pathotype III (highly aggressive), but recently added a new *A. rabiei* pathotype (pathotype IV) (Imtiaz et al., 2015). The new pathotype IV infected the “ highly resistant” genotypes ICC3996 and ICC12004. Worryingly, only low levels of resistance were identified in advanced breeding material screened against the pathotype IV isolate at the International Center for Agricultural Research in the Dry Areas (ICARDA) (Imtiaz et al., 2015). Most recently in Australia, a broad range of aggressiveness was observed including several highly pathogenic isolates (Mehmood et al., 2017). These were not associated with a host genotype or growing region, however under controlled environmental conditions, they were able to cause severe disease symptoms on moderately resistant PBA HatTrick and “resistant” Genesis 090 (Moore et al., 2016), two pillar cultivars of the Australian chickpea industry. Investigation of the physiological interactions between pathogen and host, at the pre and post penetration stages (Sambasivam et al., 2017 and Dadu et al., 2018) will lead to a better understanding of host defence mechanisms and to improved fungicide management.

Whilst the genetic components of the chickpea defence mechanisms to *A. rabiei* have been described in detail using molecular genomics and transcriptional approaches (Coram and Pang, 2005 and 2006; Millan et al., 2006; Rubiales and Fondevilla, 2012; Leo et al., 2016; Li et al., 2017), a comprehensive analysis of the physiological interactions is yet to be performed. Previous studies reported that germination of *A. rabiei* spores occurs at 12-48 hours post inoculation (hpi), followed by further elongation of the germ tube. The elongating germ tube then ramifies on the leaf surface and penetrates directly through the cuticle and the hyphae grow subcuticularly before proceeding deeper (Hohl et al., 1990; Pande et al., 2005). After penetration, hyphae grow between epidermal and parenchyma cells, and consume the inner leaf structures with symptoms visible after 3 to 5 days (Pande et al., 2005). However, an in-depth histological study of the timing of underlying responses, particularly prior to initiation of physiological and biochemical defence responses at 24 hpi, has not been reported. This is pivotal to understanding the timing and recognition of the pathogen by the host in the very early but crucial host-pathogen interaction stage (Sambasivam et al., 2017; Dadu et al., 2018) and when it may be possible to control the pathogen through prophylactic chemical applications. Thus, the aims of this study were to: 1) assess recent changes in aggressiveness within the Australian *A. rabiei* population; 2) determine early virulence responses of chickpea cultivars to highly aggressive isolates through the assessment of physiological and biochemical factors; and 3) provide evidence of potential pathogenicity groups (PGs) within the Australian *A, rabiei* population based on gross disease symptomology and microscopic physiological evidence that may also be used as descriptors to inform on cultivar longevity and fungicide management strategy.

## Materials and Methods

### Pathogenic structure and classification of highly aggressive Australian *A. rabiei* isolates into Pathogenicity Groups

#### Plant materials

Five chickpea genotypes with known disease reactions were used as a differential host set to assess the aggressiveness of the Australian *A. rabiei* population: ICC3996, Genesis 090, PBA HatTrick, PBA Seamer and Kyabra (**Table 1**). Seed was obtained from the National Chickpea Breeding Program, Tamworth, NSW, Australia. Seedlings were grown in 15 cm diameter pots containing commercial grade potting mix (Richgro premium mix), fertilized by Nitrosol, Amsgrow ® (4.5 mL/L) every two weeks and watered as required. The experiment was conducted with a complete randomised design with two replicates for each genotype × isolate combinations assessed, with five plants per pot/rep. All plants were grown and maintained in the controlled environment facility at 22 ± 2°C under 16 h/8 h day/night photoperiod with near-UV light irradiation of 350–400 nm (Supplied by Valoya LED grow lights, Finland) at Griffith University, Nathan campus, Queensland, Australia.

**Table 1.** Differential host genotypes used in the bioassay and their disease ratings to *A. rabiei* in Australia.

| Genotype | Type | Pedigrees | Resistance level | Reference |
| --- | --- | --- | --- | --- |
| **ICC3996** | Desi | ICC3996_x_ILWC184-F2 x Lasseter_x_ICC3996-F6-RIL | Resistant (R) | Nasir et al., (2000); Moore et al., 2015 |
| **PBA Seamer<sup>Ⓢ</sup>** | Desi | Evaluated (as CICA0912) and developed by PBA chickpea breeding Program from a cross between breeding line 98081-3024 and PBA HatTrick<sup>Ⓢ</sup> | Moderately resistant (MR) | Pulse Australia (2020) |
| **Genesis 090** | Kabuli | FLIP82-150C/FLIP83-48C | MR/Moderately susceptible (MS) | Pulse Australia (2018) |
| **PBA HatTrick<sup>Ⓢ</sup>** | Desi | Jimbour x ICC14903 | MS | Pulse Australia (2018) |
| **Kyabra** | Desi | AMETHYST//946-31/BARWON//LASSETER/940-26//946-31/NORWIN//8507-28H//AMETHYST//T1069/8507-28H//946-31 | Very susceptible (VS) | Pulse Australia (2018) |

#### Fungal materials, inoculation and disease assessment

201 single spored *A. rabiei* isolates were selected for phenotyping, representative of the years, regions and host genotype origins sampled within the 2016-2017 collection as described in Mehmood et al. (2017). The full list of isolates and their available passport data (place of collection, year of collection, and host genotype) is provided in the additional material (**Online source 1**).

Single spored isolates were cultured in V8 juice agar and maintained in the incubator for 14 days at 22 ± 2°C with a 12/12 h near-UV light irradiation (350–400 nm)/dark photoperiod prior to being used in the inoculation bioassay. Spore suspensions were prepared by harvesting pycnidiospores from fourteen-day-old fungal cultures by flooding with sterile water and gently disturbing the surface with a sterile glass rod. The spore suspension was filtered through cheese cloth (250 mm) to separate conidia from mycelia and inoculum concentrations were adjusted to 1’10^5^ spores/mL using a haemocytometer. Two to three drops of Tween 20 (0.02% v/v) were added per 100 mL of spore suspension as a surfactant and inoculation was carried out using the modified mini-dome technique (Chen et al., 2005, Sambasivam et al., 2017). Briefly, plants were inoculated until run-off using a 500 mL hand sprayer producing a fine mist of inoculum, and the pots were rotated during the inoculation procedure to achieve an even spread of inoculum. The differential host set plants were placed randomly in a separate solid 20 L plastic crate for inoculation with a single isolate and sealed for 24-36 hours to provide dark and high humidity conditions. At 24 hours post inoculation (hpi), the lids were removed, and plants were left until disease assessment. Plants were misted twice daily to provide conditions conducive for disease development. Disease severity of each isolate on each of the host genotypes was assessed using the qualitative 1–9 scale of Singh et al. (1981) at 21 days post inoculation (dpi) where; scores of 1 or 3 represented a low disease severity; 5 represented a moderate disease severity without significant stem infection, and 7 or 9 represented a high disease severity with stem lesions that would lead to major difficulties in transpiration, photosynthesis and possible stem breakage.

#### Detection of highly aggressive isolates and their pathogenicity grouping (PG)

Isolates were identified as highly aggressive when they produced a leaf score of at least 7 on >80%, and a stem score of at least 7 on >10%, of the number of plants assessed for each host x isolate combination. Subsequently, the highly aggressive isolates were further classified into PG 1-5 based on their ability to cause low, moderate or high disease severity independently on ICC3996, Genesis 090 and PBA HatTrick (**Table 2**).

**Table 2.** Criteria used for pathogenicity grouping of the highly aggressive isolates.

| **Pathogenicity Group** | **Description** |
| --- | --- |
| 0 | Low disease on PBA HatTrick<sup>Φ</sup>, Genesis 090 and ICC3996 |
| 1 | High disease on PBA HatTrick<sup>Φ</sup> and low disease on Genesis 090 and ICC3996 |
| 2 | High disease incidence on PBA HatTrick<sup>Φ</sup>, moderate disease on Genesis 090 and low disease on ICC3996 |
| 3 | High disease on PBA HatTrick<sup>Φ</sup>, moderate disease on Genesis 090 and moderate disease on ICC3996 |
| 4 | High disease on PBA HatTrick<sup>Φ</sup>, high disease on Genesis 090 and moderate disease on ICC3996 |
| 5 | High disease on PBA HatTrick<sup>Φ</sup>, Genesis 090 and ICC3996 |

### Physiological and biochemical differences observed among host-isolate interactions *Differences among aggressiveness traits between isolates from different Pathogenicity Groups*

#### Plant material, fungal material and inoculum preparation

Seed of genotypes ICC3996, Genesis 090, PBA HatTrick and Kyabra were grown as described above. Single spored isolates were chosen from PG 0-4 within the 2013-2015 populations (**Table 3**), as was determined based on the symptomology on the host set (Mehmood et al., 2017). The experiments were conducted in a complete randomised design with four replicates sown at each time point/isolate and one leaflet from each replicate used to assess the development of infection structures. One pot of each host genotype at each selected time point was mock inoculated with water and surfactant as a pathogen-free control.

**Table 3.** List of isolates selected for histopathology assessment with known levels of pathogenicity (Mehmood et al., 2017).

| Isolate | Collection year | Location | Host source | Aggressiveness and Pathogenicity Group (PG) |
| --- | --- | --- | --- | --- |
| **FT13092-1** | 2013 | Kingsford, South Australia (SA) | Genesis 090 | High – PG 4 |
| **FT13092-4** | 2013 | Kingsford, SA | Genesis 090 | High – PG 3 |
| **FT13092-6** | 2013 | Kingsford, SA | Genesis 090 | High – PG 2 |
| **TR6421** | 2014 | Yallaroi, New South Wales (NSW) | PBA HatTrick<sup>Ⓢ</sup> | High – PG 1 |
| **TR6408** | 2014 | Yallaroi, NSW | PBA HatTrick<sup>Ⓢ</sup> | Low – PG 0 |

#### Inoculation and sample collection

Fully expanded lower leaves from 14-day-old plants of ICC3996, Genesis 090, PBA HatTrick and Kyabra were detached separately for each isolate and time point just before inoculation (**Table 4**). The detached leaves were surface sterilised by immersing in a 70% ethanol solution for 30–60 s, rinsed with sterile water and then dried with sterile cotton wool. One leaflet from each of four replicates of each host x isolate interaction was inoculated and incubated as described by Sambasivam et al. (2017). Briefly, the leaves from each plant were placed with the abaxial surfaces facing upward in a separate petri dish lined with moist filter paper to provide high relative humidity. One drop (∼10 μL) of a spore suspension of 1×10^5^ conidia mL^−1^ was applied using a micropipette to the middle section of the abaxial surface of each detached leaf. Petri dishes were sealed with parafilm and incubated at 22 ± 2 °C in the dark until samples were collected (**Table 4**).

**Table 4.**
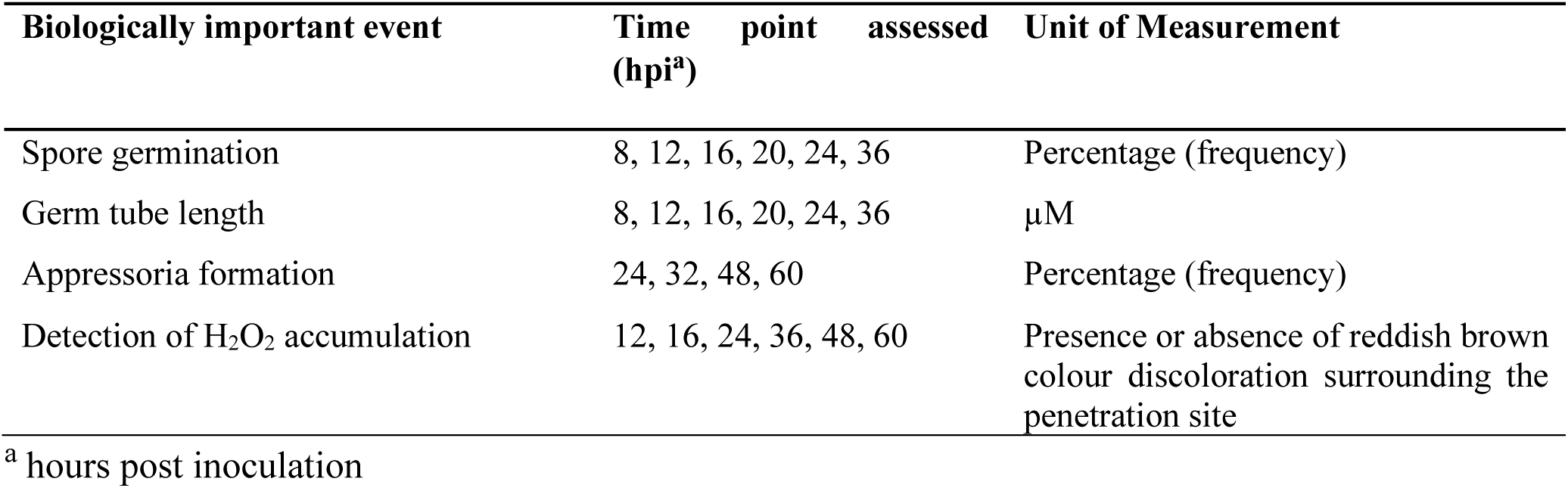
Time points selected to study each host-pathogen interaction characters.

| Biologically important event | Time point assessed (hpi<sup>a</sup>) | Unit of Measurement |
| --- | --- | --- |
| Spore germination | 8, 12, 16, 20, 24, 36 | Percentage (frequency) |
| Germ tube length | 8, 12, 16, 20, 24, 36 | μM |
| Appressoria formation | 24, 32, 48, 60 | Percentage (frequency) |
| Detection of H<sub>2</sub>O<sub>2</sub> accumulation | 12, 16, 24, 36, 48, 60 | Presence or absence of reddish brown colour discoloration surrounding the penetration site |

<sup>a</sup> hours post inoculation

#### Fixation and clearing of leaf samples

Inoculated leaf tissues were fixed and cleared as described by Achuo et al. (2004) with the following modifications: the leaf tissue was immersed in a clearing solution (absolute ethanol:glacial acetic acid; 1:2 V/V) for 24 h to remove chlorophyll. The solution was then removed using a sterile pipette and replaced with fresh clearing solution for another 12-24 h (Khan and Hsiang, 2003). The cleared leaves were stored in lactic acid/glycerol/water (1:1:1 V/V/V) at room temperature until microscopic examination.

#### Visualisation of fungal structures

Fungal structures were visualised by staining the cleared leaf tissue with lacto-phenol cotton blue (Sigma Aldrich) for 5 min or trypan blue (0.05%; Sigma Aldrich) in a phosphate buffer solution for 10 min. The stained leaf tissue was washed in 70% ethanol for a few seconds and rinsed with water in order to remove excess dye. The leaf tissue was then mounted in water on a glass microscope slide with a cover slip. The samples were examined with a Leica DMRBE light microscope fitted with a Leica DC300F digital camera. Images were captured using Leica IM50 v4 software.

#### Evaluation of interactions and fungal infection

Spore germination, germ tube length and appressoria development of each isolates was assessed on the detached leaves of each chickpea genotype throughout the experiment, between 8-60 hpi (**Table 4**).

Four independent leaves were collected at each time point (one from each of four plants) per isolate/host interaction. For each leaf, observations were taken at locations close to each of the four corners of the inoculation site. In total, 16 observations were made for each interaction at each time point. Also, a total of 100 spores were examined for each interaction at each time point (25 spores per corner) to calculate the spore germination and appressoria formation percentages. A spore was considered to be germinated when a germination tube was present, regardless of length or number of germ tubes per spore. All globular structures with a diameter larger than that of the germination tube or hyphae of origin were considered an appressoria (Dita et al. 2007). Images of three conidia were randomly selected from a single leaf from each of four replicate plants and germ tube length was measured using Leica IM50 v4 software. Four germinated germ tubes at each corner of each leaflet were assessed for appressoria development frequency (ADF) and calculated with the following formula:

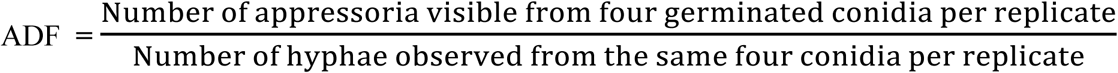

#### Histochemical detection of ROS

The production of hydrogen peroxide (H_2_O_2_) in chickpea leaves of the genotypes ICC3996, Genesis 090, PBA HatTrick and Kyabra in response to *A. rabiei* isolates FT13092-1 (PG 4), FT13092-4 (PG 3), FT13092-6 (PG 2), TR6421 (PG 1) and TR6408 was detected using the 3–3, diaminobenzidine (DAB, Sigma-Aldrich, USA) uptake method (Thordal-Christensen et al. 1997; Vanacker et al.,2000). Briefly, detached leaves of the four host genotypes were excised (five replicates per host genotype/isolate combination) into segments (1.0 × 0.5 cm) using a sterile razor blade and immediately immersed in a 1 mg ml^−1^ DAB HCl solution (pH 3.8) and left for 8 h in the dark at room temperature. After incubation, the leaf tissues were inoculated with each *A. rabiei* isolate belonging to each PG, as described previously, and samples were collected at 12, 24, 36 and 48 hpi. Isolates TR6421 and TR6408, classified as PG 1 and 0, respectively, were also sampled at 40 and 60 hpi. Samples were then fixed, cleared and stained with lacto-phenol cotton blue as previously described. Tissues were examined for H_2_O_2_ accumulation as an indication of a hypersensitive response (HR) at the locations of hyphae penetration and the surrounding cells. H_2_O_2_ was detected as a reddish-brown colouration in the DAB treated tissue and visualised under a light microscope (Leica DM750).

### Stability of aggressiveness traits among isolates from the highest aggressive Pathogenicity Groups; PG 4 and 5

#### Plant material, fungal isolates, inoculum preparation and inoculation

To test the stability of aggressiveness traits, seed of genotypes ICC3996, PBA HatTrick, PBA Seamer and Kyabra were grown as described above. Single spored highly aggressive isolates from each of PG 4 (17CUR005) and PG 5 (F17191-1) were chosen to test for differences in their aggressiveness traits on the same host set (**Table 5**). The inoculum preparation and inoculation were carried out using the methods previously descibed.

**Table 5.** Isolates selected for stability of aggressiveness traits assessment.

| Isolate | Collection year | Location | Host source | Aggressiveness and Pathogenicity Group (PG) |
| --- | --- | --- | --- | --- |
| F17191-1 | 2017 | Kingsford, South Australia | Genesis 090 | High – PG 5 |
| 17CUR005 | 2017 | Curyo, Victoria, Australia | Genesis 090 | High – PG 4 |

#### Evaluation of interactions and fungal infection

The physiological interactions, including fungal infection structures and ROS production, between each of these isolates and the differential host set genotypes, were assessed using methods previously described.

### Statistical analysis

Germination percentage, germ tube length and appressoria development frequency data were square-root transformed prior to statistical analysis and graphs were prepared with original back-transformed percentage data. Replicated measurements were averaged and the mean values with standard deviations (SD) were determined. Spore germination, germ tube length and appressoria formation data were assessed via analysis of variance (ANOVA) and further tested with least significant difference (mean values were compared using the Fisher LSD test, *P* <0.05) to determine factors that differentiated isolates between pathogenicity groups. SPSS software (IBM Corp. SPSS 2012) and Genstat (VSN International, 18^th^ edition, 2015) were used for all statistical analysis.

## Results

### Classification of highly aggressive isolates into Pathogenicity Groups (PG)

Of the 201 isolates assessed from the 2016-2017 populations, 81 were highly aggressive (40.79%) and able to cause severe disease symptoms on at least one of the three “ resistant” genotypes (**Supplementary Table 1**). When categorised into PG, 27% were assigned to PG 1, 15% to PG 2, 25% to PG 3 and 21% to PG 4. Also, 6% were classified as belonging to PG 5, which were discriminated from PG 4 isolates by their ability to cause severe disease symptoms on the ICC3996 resistance source and were first detected in 2017 (**Figure 1**).

**Figure 1.**
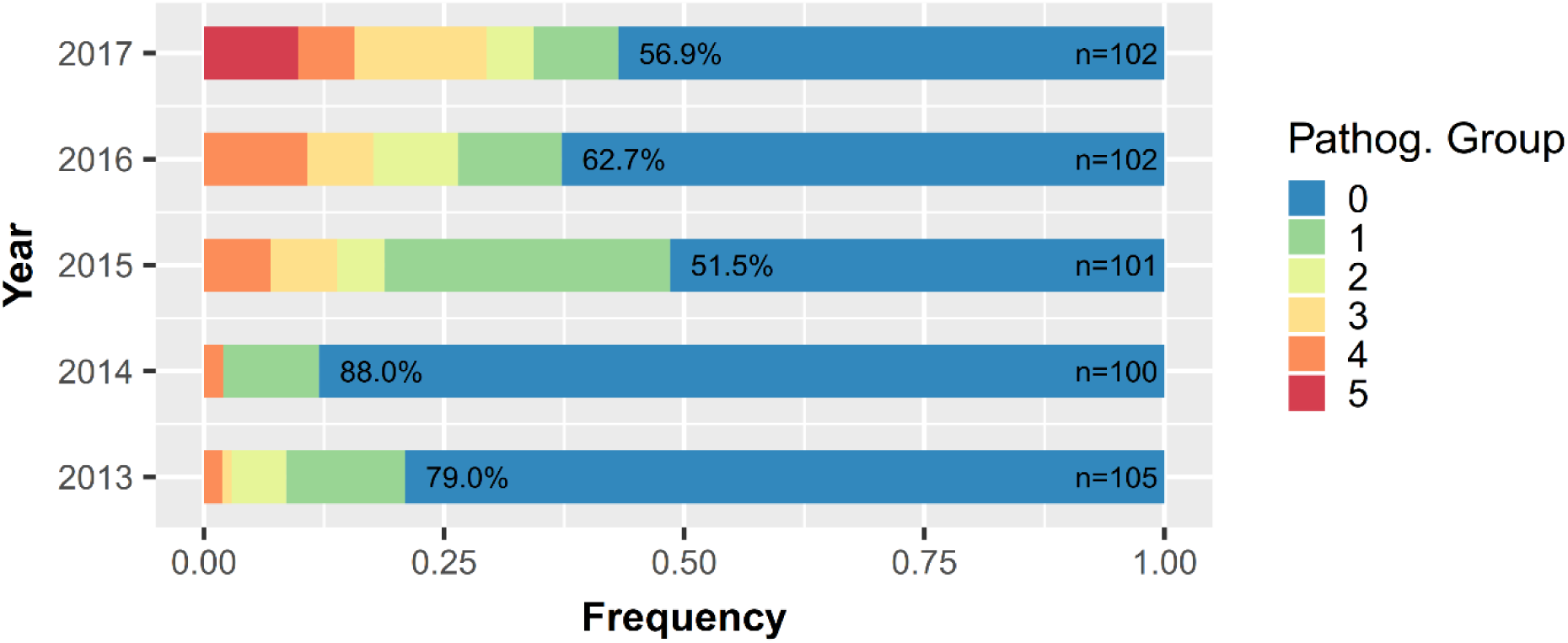
Pathogenicity Group frequencies in the 2013-2017 Australian *A. rabiei* populations.

### Physiological and/or biochemical differences among the host-isolate interactions *Differences among aggressiveness traits between isolates from different Pathogenicity Groups*

#### Spore germination percentage

Spores of isolate FT13092-6 (PG 2) germinated significantly faster than any other isolate on all host cultivars, with germ tubes visible as early as 12 hpi. This was followed by a rapid increase in spore germination percentages at 16-24 hpi on cultivars Kyabra, PBA HatTrick and Genesis 090, peaking at 36 hpi, at which 100±2.23%, 98.98±3.67% and 91.20±1.34% of the spores had germinated on the three hosts, respectively. This was significantly higher than any other isolate (*P*≤0.05, **Figure 4**). A similar trend was observed for isolate FT13092-4 (PG 3), although a lower maximum spore germination percentage of approximately 70% at 36 hpi was reached on the same hosts. In contrast, isolates TR6421 (PG and TR6408 (PG 0) showed significantly lower germination rates (<10%) until 20 hpi, then increasing to a maximum of approximately 50% on Kyabra and PBA HatTrick and this was lower on Genesis 090 (22% and 35%, respectively). Germination of isolate FT13092-1 (PG 4) spores was somewhat intermediate, with lower overall germination rates, reaching a maximum of 60%. These trends were also observed on the resistant cultivar ICC3996, however, the overall spore germination rate on this host was substantially lower for all of the isolates assessed, with a maximum of just 50% for isolate FT13092-6 at 36 hpi. Isolates FT13092-1 and FT13092-4 showed similar responses on ICC3996, beginning germinating at 16 hpi and reaching a maximum spore germination rate of approximately 30%. Meanwhile both TR6421 and TR6408 showed no spore germination until 24 hpi and reached a significantly lower maximum of approximately 15% after 36 hpi (*P*≤0.05, **Figure 2**). This was significantly higher than for FT13092-1 (PG 4) on all four host genotypes (*P*≤0.05).

**Figure 2.**
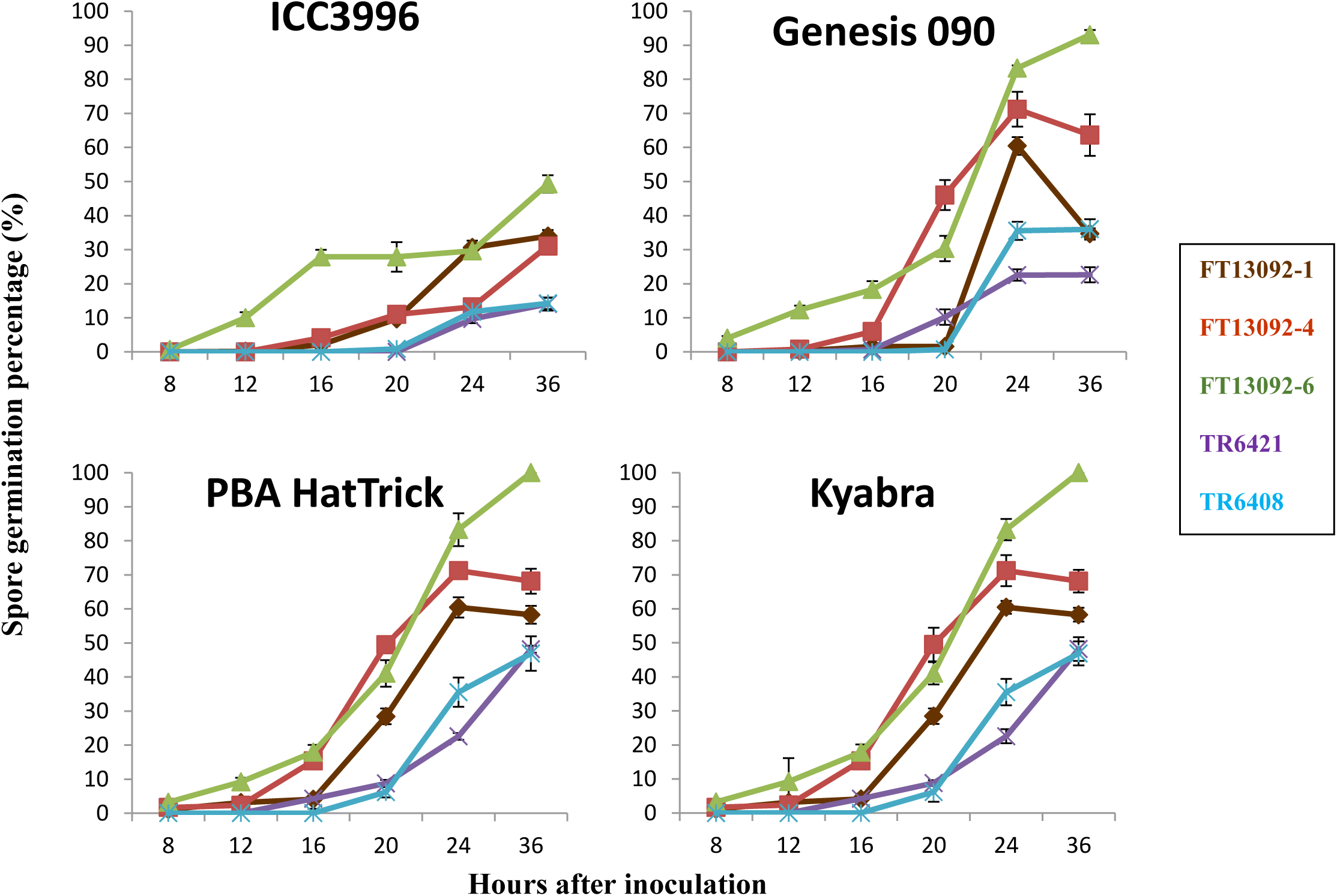
Spore germination percentage of each isolate on each host genotype over time after inoculation.

**Figure 3.**
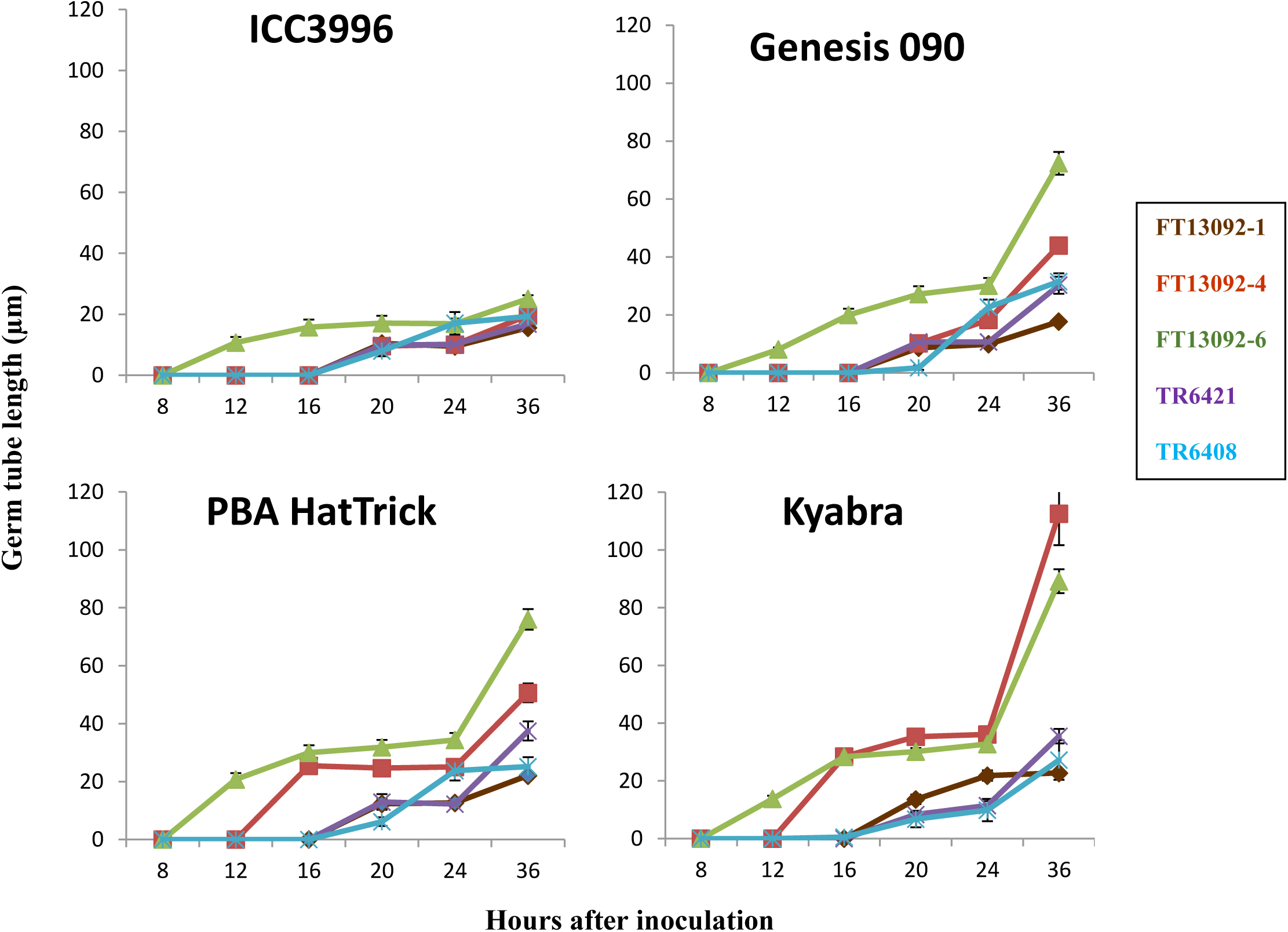
Germ tube length of each isolate on each host genotype over time after inoculation.

**Figure 4.**
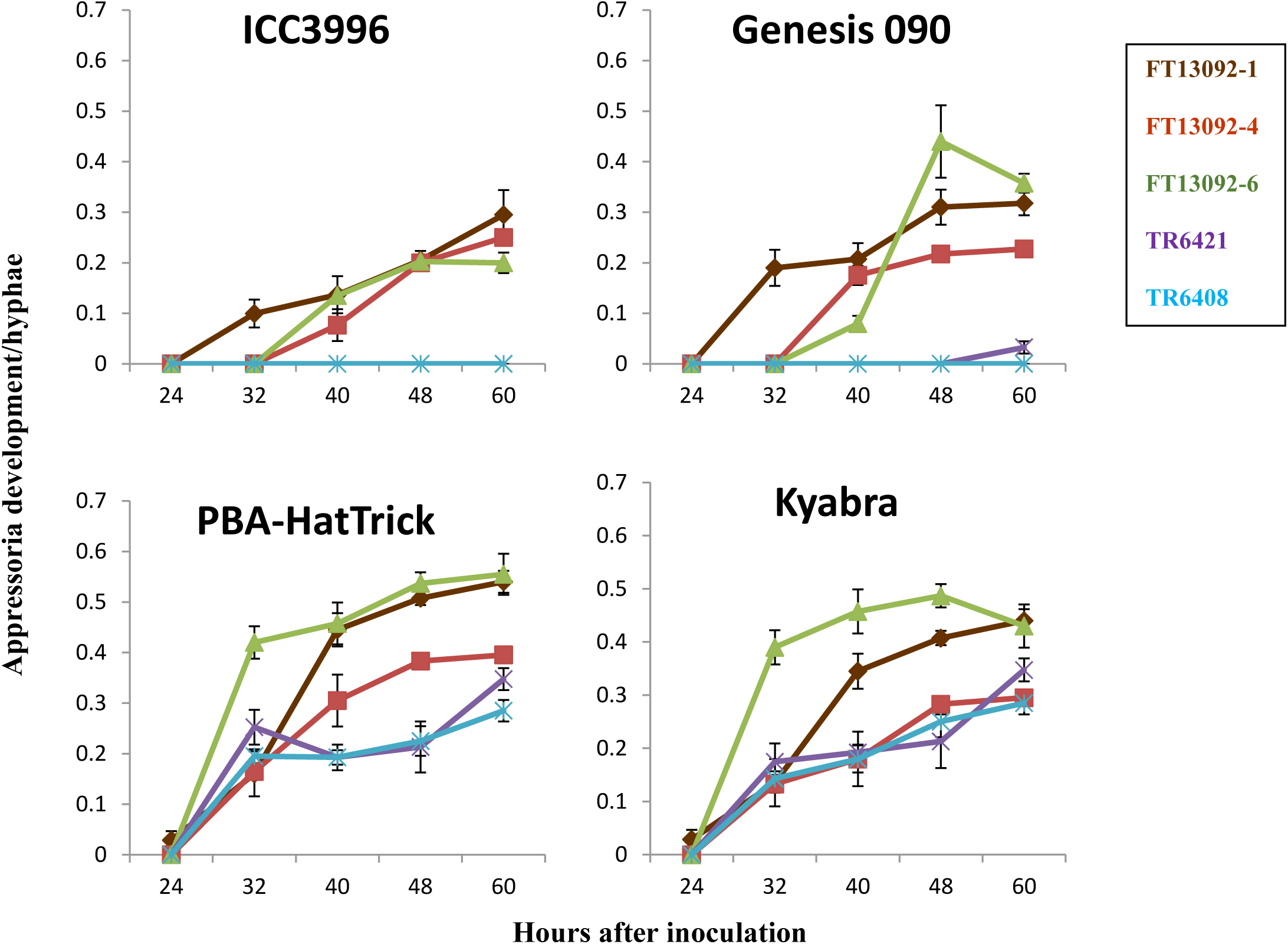
Appressoria development of each isolate on each host genotype over time after inoculation.

#### Germ tube length

Isolate TR6408 (PG 0), exhibited a significantly shorter germ tube length than all highly aggressive isolates on all genotypes assessed (*P*≤ 0.05). Meanwhile, the germ tube lengths for all isolates assessed were significantly shorter on ICC3996 than on all other hosts assessed (*P*≤0.05). The highly aggressive isolate, FT13092-6 had a significantly longer germ tube length on all hosts than the low aggressive isolate, TR6408 (*P*≤0.05). A significantly shorter germ tube length was observed for isolate FT13092-6 on ICC3996 (25.08±2.76 μm) than on Genesis 090 (72.08±3.91 μm), PBA HatTrick (76.08±3.56 μm) and Kyabra (89.16±4.15 μm) at 36 hpi (*P*≤0.05) (**Figure 3**).

#### Appressoria development frequency

The appressoria development frequency of isolate FT13092-6 (PG 2) was significantly higher on Kyabra, PBA HatTrick and Genesis 090 than on ICC3996 (*P*≤0.05) at all time points assessed (**Figure 4**). The overall appressoria frequency was highest on Kyabra followed by on PBA HatTrick and Genesis 090 across all time points and isolates except at 32 hpi and 40 hpi where it was higher on PBA HatTrick and Genesis 090 respectively. Only a few appressoria were observed on ICC3996. With the exception of isolate FT13092-1, which formed appressoria at 36 hpi, the appressoria development was generally delayed until 40 hpi on ICC3996 and Genesis 090. In contrast, all five isolates had developed appressoria on Kyabra and PBA HatTrick by 32 hpi.

At 60 hpi, the appressoria frequency was significantly lower on ICC3996 and Genesis 090 (*P*≤0.05) for two highly aggressive isolates, FT13092-6 (0.35±0.020 and 0.20±0.018, respectively) and FT13092-1 (0.31±0.048 and 0.29±0.023, respectively). However, this was higher on PBA HatTrick and Kyabra i.e. FT13092-6 (0.55±0.037 and 0.44±0.025, respectively) and FT13092-1 (0.54±0.040 and 0.43±0.021, respectively). A similar observation was made for isolates FT13092-4 and TR6421. Isolate TR6408 produced significantly more appressoria on PBA HatTrick (0.28±0.025) and Kyabra (0.29±0.021) (P<0.05). Interestingly, TR6408 did not develop any appressoria on Genesis 090 or ICC3996 by 60 hpi (**Figure 4**).

#### Histochemical detection of reactive oxygen species (ROS)

Hydrogen peroxide (H_2_O_2_), as a marker of an hypersensitive response (HR), was detected at appressoria locations of FT13092-1 (PG 4), FT13092-4 (PG 3), and FT13092-6 (PG on ICC3996 and Genesis 090, and as early as 36 hpi (**Figure 5**). Similar but less frequent observations were seen on PBA HatTrick and Kyabra in response to FT13092-6 (PG 2), between 36-40 hpi and with no HR was observed on PBA HatTrick and Kyabra in response to FT13092-1 or FT13092-4 (**Figure 5**).

**Figure 5.**
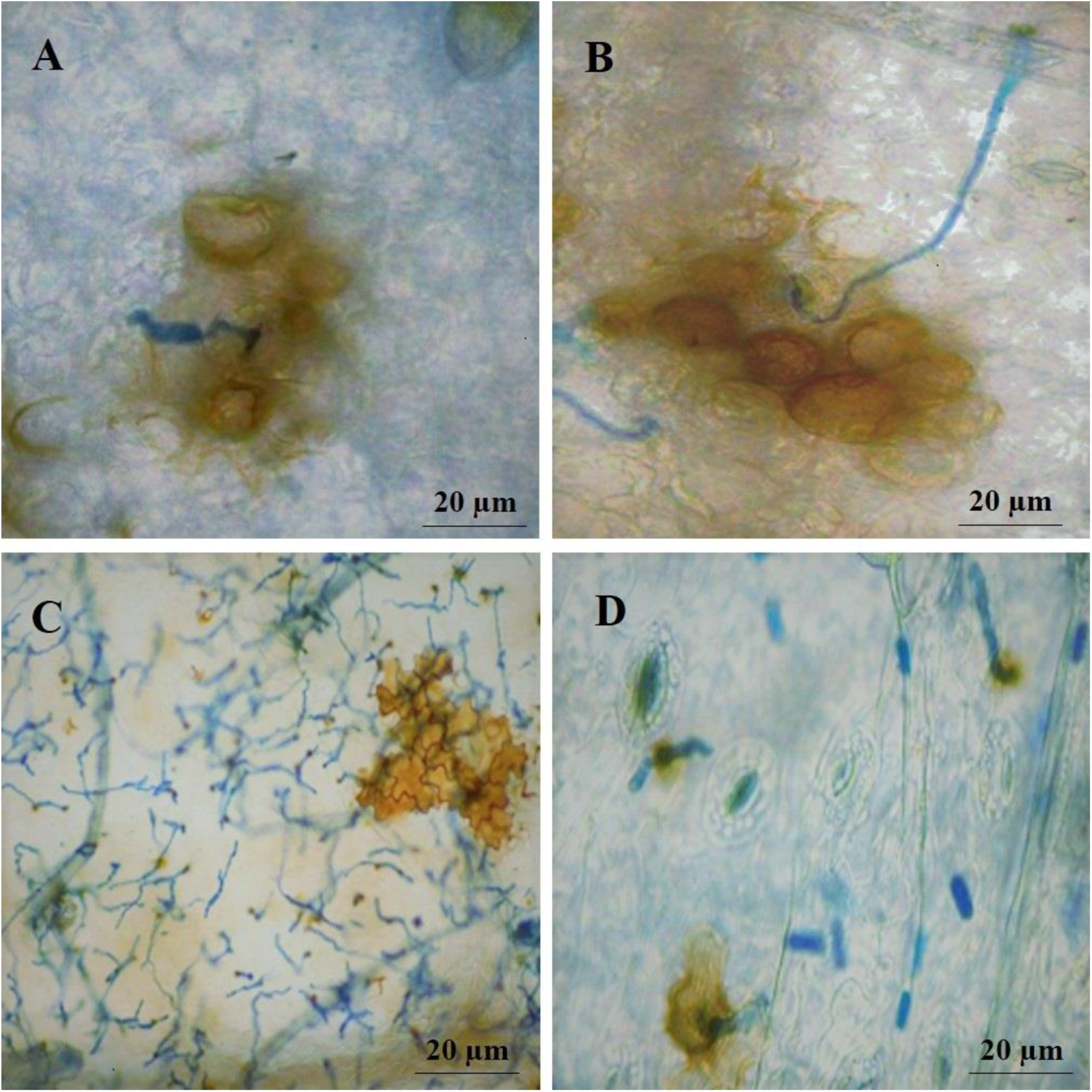
The hypersensitive response observed following inoculation with isolate FT13092-1 in **A**) ICC3996 and **B**) Genesis 090, at 36 hpi; **C**) PBA HatTrick and **D**) Kyabra, at 60 hpi.

By 36 hpi on ICC3996 and Genesis 090, H_2_O_2_ had accumulated at the penetration sites and was observed in the surrounding epidermal cells. In addition to H_2_O_2_ production, other typical characteristics of an HR were observed such as a thickening of cell walls (**Figure 5A-B**). These were not observed in PBA HatTrick and Kyabra until 60 hpi (**Figure 5C-D**). Interestingly, an HR was not observed on ICC3996 and Genesis 090 in response to TR6421 (PG 1) until 60 hpi. Similarly, the low aggressive isolate TR6408 did not instigate H_2_O_2_ accumulation until 60 hpi on PBA HatTrick. The frequency of this observation was far lower than that observed for the highly aggressive isolates and was also not observed on ICC3996 and Genesis 090 until 80 hpi.

### Stability of aggressiveness traits among isolates from the highest aggressive Pathogenicity Groups; PG 4 and 5

#### Spore germination percentage

Isolates F17191-1 (PG 5) and 17CUR005 (PG 4) germinated at 6 hpi on Kyabra, much earlier than on the other genotypes assessed. Also, a far lower percentage of spore germination was observed on PBA Seamer and ICC3996 for both isolates than on the other host genotypes (**Figure 6**). At 24 hpi, germination was significantly different on each of the hosts for both isolates assessed (P<0.05). This was highest on Kyabra with 62.67% (F17191-1) and 66.87% (17CUR005) germination at 24 hpi. Spore germination percentage was lower on PBA Seamer (42.07%; F17191-1 and 44.17%; 17CUR005), ICC3996 (44%; F17191-1 and 56.50%; 17CUR005) and PBA HatTrick (46%; F17191-1 and 49.33%;17CUR005). Spore germination percentage on PBA Seamer, ICC3996 and PBA HatTrick was lower when compared with Kyabra at all time points observed for both isolates, in agreement with the level of host resistance revealed through disease reaction phenotyping. The lower germination was observed on both PBA Seamer and ICC3996 until the final observation at 36 hpi (**Figure 6**).

**Figure 6.**
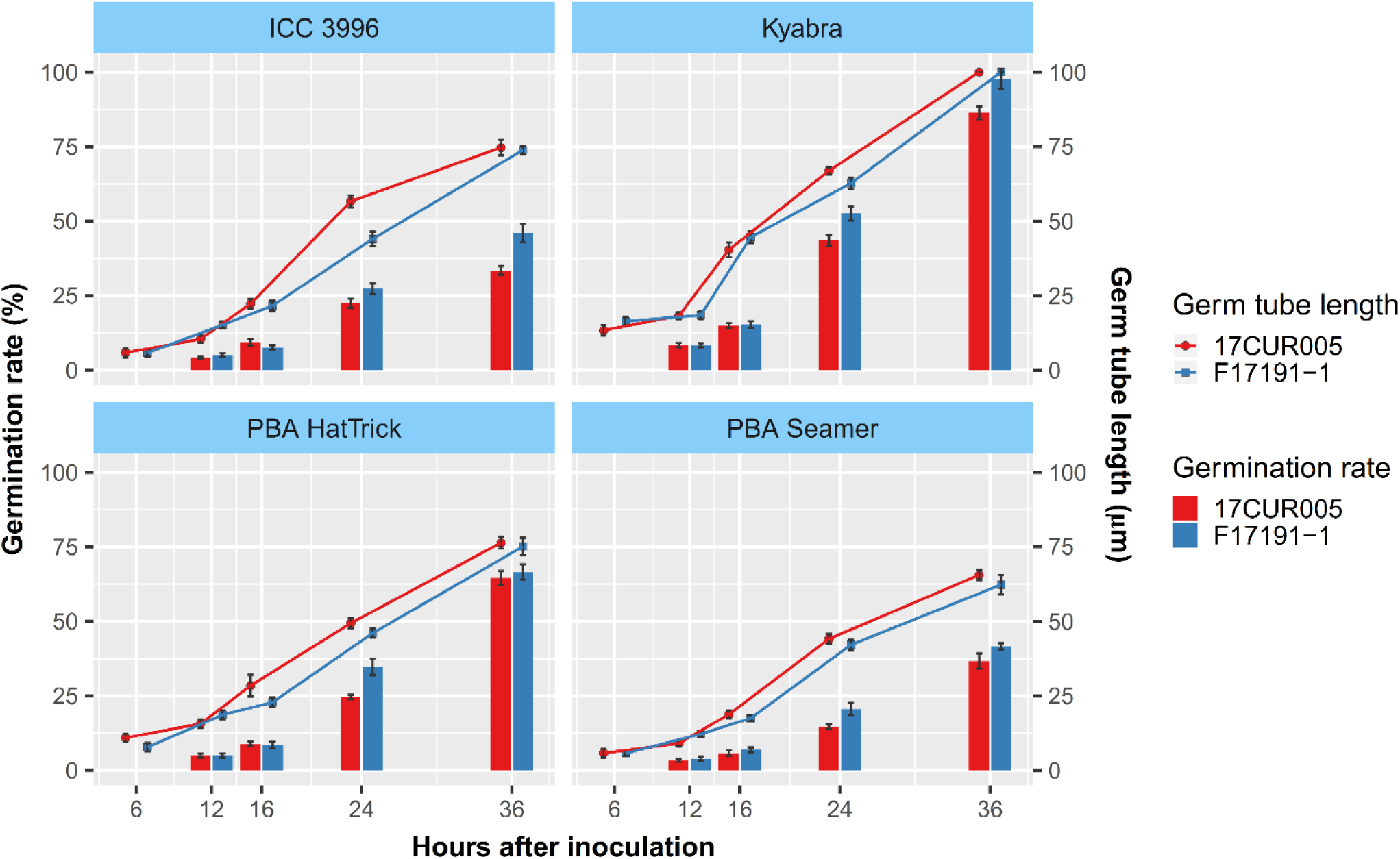
Spore germination (%) and germ tube length (µm) of isolates F17191-1 and 17CUR005 on each of the four host genotypes over the time points assessed.

#### Germ tube length

Isolates F17191-1 and 17CUR005 developed much shorter germ tubes on ICC3996, PBA Seamer and PBA HatTrick and appeared to penetrate much faster than when grown on Kyabra. At 24 hpi, germ tubes were significantly shorter on PBA Seamer (20.59 µm; F17191-1 and 14.59 µm; 17CUR005) than on ICC3996 (27.30 µm; F17191-1 and 23.45 µm; 17CUR005), PBA HatTrick (34.74 µm; F17191-1 and 24.60 µm; 17CUR005) and far shorter than on Kyabra (52.61 µm; F17191-1 and 43.48 µm; 17CUR005) (*P*<0.05). The length differences persisted until final assessment when germ tubes of both isolates were significantly shorter on PBA Seamer (41.58 µm; F17191-1 and 36.65 µm; 17CUR005) than on ICC3996 (46.01 µm; F17191-1 and 33.44 µm; 17CUR005) and PBA HatTrick (66.50 µm; F17191-1 and 64.48 µm; 17CUR005) and were far shorter than on Kyabra (97.75 µm; F17191-1 and 86.28 µm; 17CUR005) at 36 hpi (*P*<0.05) (**Figure 6**).

#### Histochemical detection of ROS generation in response to highly aggressive isolates

Similar but less frequent observations related to an HR were observed on PBA HatTrick in response to either F17191-1 or 17CUR005 at 36 hpi and 48 hpi and no HR was observed on Kyabra in response to either isolate. At 48 hpi and 60 hpi on ICC3996 and PBA Seamer, H_2_O_2_ accumulated strongly at the penetration sites and in the surrounding epidermal cells in response to both isolates (**Figure 7A-B**). In addition to H_2_O_2_ production, other typical scharacteristics of an HR were observed such as a thickening of cell walls in PBA HatTrick (**Figure 7C**). These responses were not observed in Kyabra until 60 hpi (**Figure 7D**) for either isolate. Notably, isolate 17CUR005 triggered H_2_O_2_ production at 48 hpi in PBA Seamer but this was far less than observed in ICC3996 or PBA HatTrick before 60 hpi.

**Figure 7.**
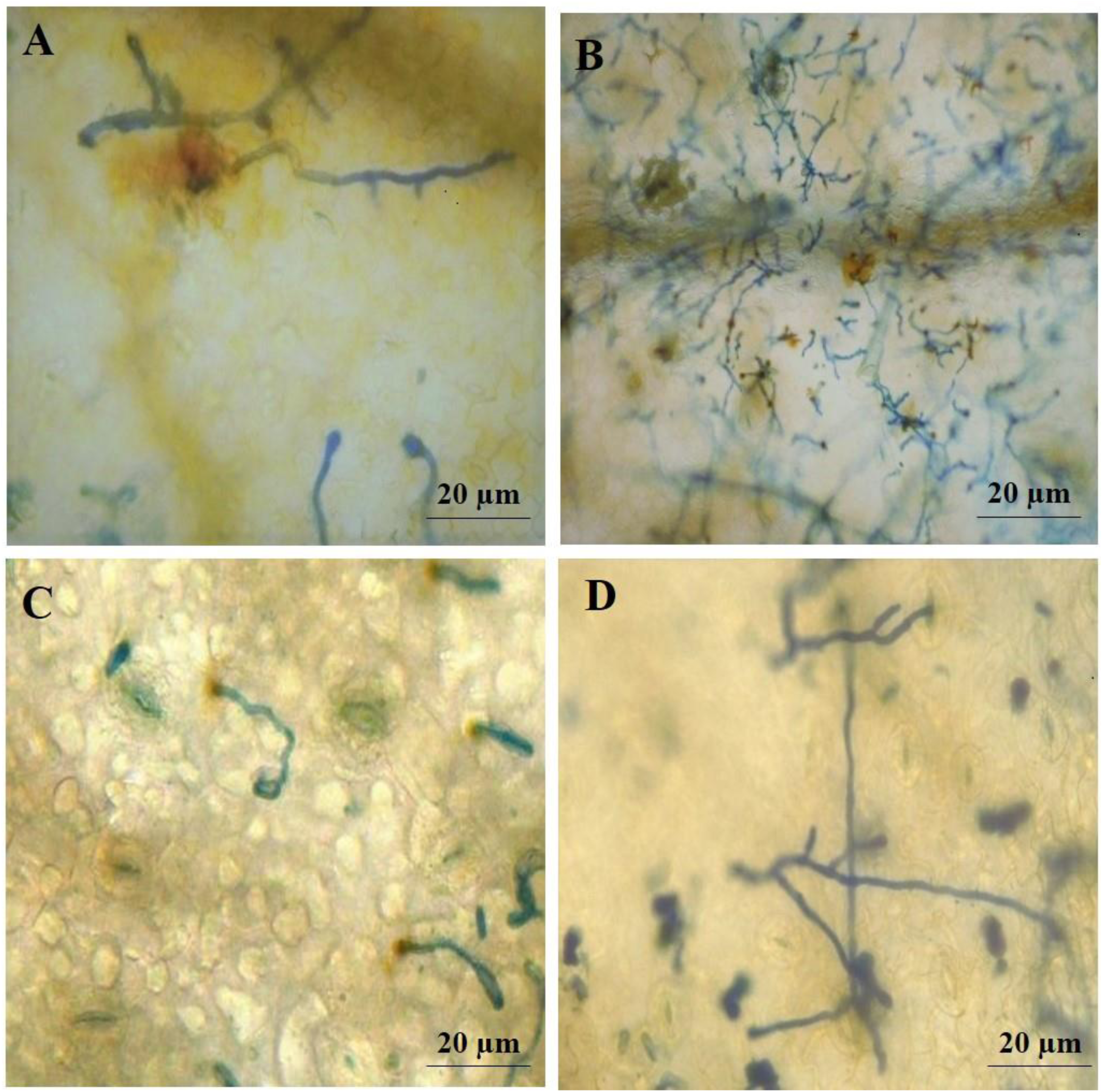
The hypersensitive response observed for highly aggressive isolate (F17191-1; PG 5) in: **A**) PBA Seamer at 36 hpi **B**) ICC3996 at 48 hpi, and **C**) PBA HatTrick at 48 hpi and **D**) No HR observed in Kyabra until 60 hpi.

## Discussion

### Classification of highly aggressive isolates into Pathogenicity Group (PG)

Understanding the aggressiveness of a pathogen is valuable in improving the effectiveness of management practices, particularly in the strategic use of cultivar resistance and timing and type of fungicide application against a foliar-infecting fungus. Accordingly, assessing the recent population of Australian *A. rabiei* isolates on a chickpea differential host set demonstrated a natural diversity in aggressiveness that had similarly been detected over prior time periods and elsewhere (Atik et al., 2013; Benzohra et al., 2014; Mahiout et al., 2015; Mehmood et al., 2017; Baite and Dubey, 2018). Additionally, the most aggressive isolates were identified for use in future selective breeding strategies.

The subsequent classification of the highly aggressive isolates into pathogenicity groups based on their ability to cause disease on the best resistance sources was backed up with significant differences observed in physical and biochemical factors among the select few isolates assessed. Meanwhile, over the past few years, a significant increase in frequencies of isolates able to cause severe damage on formally “resistant” cultivars has occurred. In particular, the proportion of isolates that are highly aggressive on PBA HatTrick has risen from 18% in 2013 to 68% in 2017. Also, more isolates able to cause moderate to severe disease on Genesis 090 (31.68% in 2016 to 50% in 2017), a major cultivar grown in southern Australia, have been detected. Most recently, increased aggressiveness within the Australian *A. rabiei* population was highlighted by the more recent detection of highly aggressive Pathogenicity Group 5 isolates, which were able to overcome resistance in ICC 3996, one of the key sources of resistance in the chickpea breeding program, and in field trials (Moore et al., 2016) and commercial crops of PBA Seamer, a new variety that when released in 2016 was done so with an Ascochyta rating of MR (Pulse Australia, 2020).

Previously, increased susceptibility to *A. rabiei* was observed in PBA HatTrick (Moore et al., 2016; Mehmood et al., 2017), indicating potential increase in isolate aggressiveness as a selective response. To that point, Pathogenicity Group 4 isolates were the most aggressive isolates detected, able to cause high disease severity on PBA HatTrick and Genesis 090 and only able to cause moderate disease severity on ICC3996 (Mehmood et al., 2017). Consequently, erosion of resistance in host genotypes that contain alleles from the same resistance source has become a high risk within the Australian resistance breeding program, particularly noting that PBA Seamer and PBA HatTrick are progeny of crosses containing ICC3996 as the resistance donor parent. It was noted that one isolate (F17076-3) caused high disease severity on ICC3996 but not on PBA Seamer, potentially indicating that PBA Seamer does contain a selection of resistance alleles providing resistance to a sub-population of the scurrent highly aggressive isolates.

The most aggressive and commonly detected isolates are required for selective breeding and, potentially, informing disease management strategies, particularly if differential factors underpinning aggressiveness can be dissected. It is highly likely more isolates capable of causing severe damage on PBA Seamer and other subsequently released ‘resistant” varieties will evolve. The increase in aggressiveness seen over the last few years has likely occurred through “pathogen adaptation” - in other words, the fungus has evolved to produce more highly aggressive isolates due to selective pressures being applied through environmental factors and farming practices.

### Differences among aggressiveness traits between isolates from different Pathogenicity Groups

Major difference in pre and post penetration behaviours between the isolates from different pathogenicity groups included in this study were observed on leaves of the susceptible genotype Kyabra, the moderately resistant genotype PBA HatTrick and the previously resistant genotypes Genesis 090 and ICC 3996.

Initial differences were observed in spore germination at the first stage of the infection process. Conidia of all isolates germinated by 6 hpi under the conditions of the controlled environment study, irrespective of isolate/ host genotype combination, as was previously observed in other studies (Pande et al., 2005). In terms of guiding informed chemical management, spores that take longer to germinate may also be exposed longer to surface/topical/frontline/prophylactic fungicides and hence be easier to manage with these chemicals (Slawecki et al., 2002). Accordingly, Peres et al. (2010) found that Captan (protectant fungicide) was effective against *Colletotricum acutatum* when applied at 4-24 hpi but was ineffective after 24 hpi (Curry et al., 2002). Meanwhile, the delay in spore germination on the previously resistant host may suggest a purposeful reorganisation of pathogen-activated molecular patterns switching, to delay recognition and hence delay the onset of host defence responses, as identified with *A. lentis* on lentil (Ford et al., 2017). In a study conducted on wheat leaves, spore germination percentage of *Puccinia triticina* was higher on susceptible than on resistant cultivars, indicating that the resistant response was initiated right at the start of the host-pathogen interaction (Wesp-Guterres et al., 2013). In a similar study on lentil, significant differences were found in spore germination among low and highly aggressive isolates on resistant and susceptible cultivars to *Ascochyta lentis* (Sambasivam et al., 2017). Also, isolates with higher germination percentages may produce more appressoria and this most likely lead to a higher disease incidence on the host.

Potentially, in the current study, the highly aggressive isolates of *A. rabiei* induced a different expression profile of defence genes and earlier than the low aggressive isolates, adding weight to the theory that highly aggressive isolates can recognise the host faster (Kavousi et al., 2009; Pitzschke et al., 2009; Leo et al., 2016). In addition, once recognised, defence compounds excreted by the plant may interfere with spore germination (Prats et al., 2007; Sánchez-Martín et al., 2012). Indeed, *A. rabiei* phytoalexin production was previously positively correlated with aggressiveness and considered an important factor in host-pathogen interaction (Jayakumar et al., 2005). Meanwhile, potential absence or a comparatively lower level of specific phytoalexins in susceptible hosts may result in higher spore germination frequency. A comprehensive, and potentially tissue-specific, comparative transcriptomics study of the resistant and susceptible responses may undercover the key genetic components of these early defence mechanisms.

Meanwhile, the germ tube length varied significantly among low and highly aggressive isolates on the differential host set included in this study, indicative of differential specific host defence responses and perhaps influenced by isolate fitness. Potentially, elicitors released by the low aggressive isolates, TR6408 and TR6421, are more rapidly recognised by the resistant hosts, ICC3996 and Genesis 090, than the susceptible host, Kyabra. This may have subsequently led to the production of antimicrobial compounds and their delivery to the leaf surface to inhibit germ tube elongation. Compounds such as flavonoids, phenols, isoflavnoids, isocoumarins, poluenes, and furanoyerpenoids are commonly produced by plants to inhibit pathogenic fungal growth at early infection stages (Kuc, 1992; Lattanzio et al., 2006). A previous study of the defence responses in chickpea to *A. rabiei* revealed induction of cross-linking of cell walls mediated by hydrogen peroxide, production of pathogenesis-related (PR) proteins (chitinase, β-1,3-glucanase, and thaumatin-like proteins), and accumulation of phytoalexins (Jayakumar et al., 2005). Coram and Pang (2006) identified several genes potentially involved in PR protein production to *A. rabiei* including proline rich protein, SNAKIN2 antimicrobial peptide, disease resistance response protein and polymorphic antigen membrane protein, which might be potentially involved in retardation of germ tube length on ICC3996 and Genesis 090.

Meanwhile, appressoria formation enables mechanical penetration by *A. rabiei*, providing an added advantage to its ability to invade through natural openings (Ilarslan and Dolar, 2002). In this study, the highly aggressive isolates, FT13092-1 (PG 4), FT13092-4 (PG 3), formed appressoria in greater numbers and faster than the low aggressive isolates on both the resistant and susceptible hosts. This indicated the efficient way in which the pathogen was able to invade quickly into the host epidermal tissues, potentially fast enough to avoid the early molecular and biochemical host defences. In a similar study on lentil, highly aggressive isolates of *A. lentis* produced a larger number of appressoria than the lower aggressive isolate (Sambasivam et al., 2017). In the current study, the least aggressive isolate TR6408 did not develop appressoria even at 60 hpi on ICC3996.

Multiple factors in plants may contribute to low frequency and delayed production of appressoria including early recognition by the host and the potential production of antifungal compounds which might relate to timing of expression of one or several genes. For example, Talbot et al. (1993) found that during appressoria formation by the rice blast fungus, *Magnaporthe grisea* on rice, the MPG1 transcript increased in resistant plants up to 60-fold and decreased the appressoria formation. Also, appressoria formation was related to expression of three *M. grisea* genes involved in mitogen-activated protein kinase (MAPK) cascades (Choi and Dean, 1997; Dean et al., 2005). The low and delayed production of appresoria in *A. rabiei* might relate to internal or external factors including remodelling of actin cytoskeleton, arbitrated by septin GTPases, and swift cell wall differentiation and regulated by external factors like perception of plant cell surface components and starvation stress (Ryder and Talbot, 2015). Again, a tissue targeted transcriptomics approach, to assess the reaction of the host in the tissues adjacent to the fungal appressoria will likely uncover isolate-host specific defence response differences.

Recognition by the host was evident during the early stage of infection where attempts by *A. rabiei* to enter the host cells triggered an HR and the release of ROS. The distinct difference in ROS species production between the compatible and incompatible interactions at the early stages of the *A. rabiei* infection paralleled differences in pathogenicity between low and highly aggressive isolates. Strong accumulation of H_2_O_2_ was detected beneath appressoria of FT13092-1 (PG 4), FT13092-4 (PG 3), and FT13092-6 (PG 2) on ICC3996 and Genesis 090 as early as 36 hpi. It appeared that once the infection structure determined a suitable penetration site, the release of cell wall degrading enzymes and/or secretion of toxins was triggered. Similar findings were observed by Sambasivam et al. (2017) and Dadu et al. (2018), who identified earlier and stronger H_2_O_2_ accumulation in a resistant genotype compared to in a moderately resistant and susceptible genotype in the lentil/*A. lentis* pathosystem. Apart from direct toxicity to the invading pathogen, accumulation of H_2_O_2_ in the epidermal cells may lead to signalling of downstream defence mechanisms (Lin et al., 2005; Hancock et al., 2007), including the hypersensitive reaction (Lam, 2004).

### Stability of aggressiveness traits among highly aggressive isolates from PG 4 and 5

No significant differences were observed between the two highly aggressive isolates assessed in depth although the rate of formation of infection hyphae varied on the genotypes tested. Similar observations were recorded with *A. lentis* on lentil/ (Sambasivam et al., 2017; Dadu et al., 2018) and *Mycosphaerella pinodes*/*Medicago truncatula* (Suziki et al., 2017) pathosystems. The ability of spores to rapidly germinate and grow germ tubes on PBA HatTrick and Kyabra may be the initial indication of defence reduction, compared to ICC3996 and PBA Seamer. This suggests that the resistant genotypes were potentially able to recognise the pathogen faster and trigger an array of defence mechanisms. Most notable was the increased rate of germination and germ tube growth that occurred from 16-24 hpi onwards especially in Kyabra and PBA HatTrick but not PBA Seamer and ICC3996. Potentially, this may indicate a critical defence point and may demonstrate differences in the defence mechanisms among these genotypes, possibly involving an inability of the Kyabra genotype to create an inhospitable environment for the fungus on the exterior phyloplane. Instead, PBA Seamer and ICC3996 were potentially able to recognise the pathogen faster and muster antifungal compounds to the environment causing the fungus to penetrate more slowly. Another interesting feature of the resistant interaction was the observation of reduced hyphae length in or around the epidermal cells which was not observed on the susceptible genotype Kyabra. This suggested the hyphae were aberrated, which could be a structural defence response (Suzuki et al., 2017). During the penetration process, the host cell wall became degraded and/or swollen near the infection hyphae. This degradation and swelling of the cell wall was likely associated with cell wall-degrading enzymes secreted by the growing hyphae. Similar findings were reported in the *M. pinodes*/*M. truncatula* pathosystem (Suzuki et al., 2017). Similarly, the higher H_2_O_2_ production observed on PBA Seamer and ICC3996 was a well characterised defence response that causes deposition of lignin for physical strengthening and signalling of downstream defence mechanisms.

## Conclusion

A broad and natural range of early aggressiveness exists among Australian isolates of s*A. rabiei*. We believe the aggressiveness of a particular isolate is directly related to the commercial chickpea varieties they have been exposed to and the genotypes on which they are assessed. We also propose that the development of aggressiveness is influenced by the ability of the host to slow fungal growth externally and following penetration within the epidermal tissues. A more aggressive and potentially successful isolate is able to move faster into the host to evade external defences.

The physiological and biochemical analysis of the current highly aggressive Pathogenicity Group isolates provides evidence of differences in aggressive traits on the differential set, suggesting the general defence responses were expressed more rapidly and extensively in the resistance host compared to the susceptible host. There is an urgent need to identify novel resistance sources to these highly aggressive isolates to improve the sustainability of the chickpea industry and to assess the potential time to loss of resistance within the most recently released cultivars. Simultaneously, there is a need to carefully target future fungicide strategies towards effective management of the specific isolate population and at the most appropriate biological stage of interaction with the host. This will include using a prophylactic surface-acting fungicide, when the fungus has landed and attached to the host on the external plant surface, prior to penetration and using a systemic internal-acting fungicide once the fungus has invaded into the tissues. The information generated in this study may be used to develop cultivar-specific fungicide regimes based on knowledge of the local pathogen population and the chickpea cultivar grown.

## Acknowledgement

The *A. rabiei* isolates were kindly supplied by DEDJETR, Horsham, Victoria; SARDI, South Australia; Department of Primary Industries, Tamworth, New South Wales (NSW) and Curtin University, Western Australia. Gail Chiplin from DPI, Tamworth, NSW and Marzena Krysinska-Kaczmarek from SARDI provided technical assitance. Seed material was kindly supplied by Kristy Hobson, DPI, Tamworth, NSW.

